# MS2DeepScore - a novel deep learning similarity measure for mass fragmentation spectrum comparisons

**DOI:** 10.1101/2021.04.18.440324

**Authors:** Florian Huber, Sven van der Burg, Justin J.J. van der Hooft, Lars Ridder

## Abstract

Mass spectrometry data is one of the key sources of information in many workflows in medicine and across the life sciences. Mass fragmentation spectra are considered characteristic signatures of the chemical compound they originate from, yet the chemical structure itself usually cannot be easily deduced from the spectrum. Often, spectral similarity measures are used as a proxy for structural similarity but this approach is strongly limited by a generally poor correlation between both metrics.

Here, we propose MS2DeepScore: a novel Siamese neural network to predict the structural similarity between two chemical structures solely based on their MS/MS fragmentation spectra. Using a cleaned dataset of >100,000 mass spectra of about 15,000 unique known compounds, MS2DeepScore learns to predict structural similarity scores for spectrum pairs with high accuracy. In addition, sampling different model varieties through Monte-Carlo Dropout is used to further improve the predictions and assess the model’s prediction uncertainty. On 3,600 spectra of 500 unseen compounds, MS2DeepScore is able to identify highly-reliable structural matches and predicts Tanimoto scores with a root mean squared error of about 0.15. The prediction uncertainty estimate can be used to select a subset of predictions with a root mean squared error of about 0.1. We demonstrate that MS2DeepScore outperforms classical spectral similarity measures in retrieving chemically related compound pairs from large mass spectral datasets, thereby illustrating its potential for spectral library matching. Finally, MS2DeepScore can also be used to create chemically meaningful mass spectral embeddings that could be used to cluster large numbers of spectra. Added to the recently introduced unsupervised Spec2Vec metric, we believe that machine learning-supported mass spectral similarity metrics have great potential for a range of metabolomics data processing pipelines.

## Introduction

In the rapidly growing field of metabolomics, mass spectrometry fragmentation approaches are a key source of information to chemically characterize molecules detected in mass spectrometry-based metabolomics datasets. Mass fragmentation (MS/MS also called MS2) spectra are created through the fragmentation of molecules in the mass spectrometer and consist of peaks that reflect the mass over charge (m/z) position of the resulting mass fragments. The peak intensities are reflective of the likelihood various fragmentation paths occur for the fragmented molecule. One of the core challenges in the metabolomics field is to link MS/MS spectra to the chemical structure of the fragmented metabolite. Over the last years, many computational tools have been developed to help with annotating MS/MS data ^1^. In many workflows that aim at extracting chemical information from MS/MS spectra, automated quantitative comparisons between pairs of spectra play a crucial role. Such comparisons are used to match unknown spectra to library spectra, i.e., spectra with known or reliably annotated structures ^2^, or to learn from networks (or graphs) built based on mass spectral similarity scores ^3,4^.

One key limitation in many approaches using mass spectral similarities is that often the main interest is not the degree of similarity between two spectra, but the *structural similarity* between the fragmented chemical compounds ^4–6^. There is no single absolute measure to determine such chemical structure relatedness, but in practice the structural similarity between molecules is frequently computed from molecular fingerprints: vectors that describe the presence/absence of many structural features in the molecule. Structural similarity is typically derived from molecular fingerprints and used as a central measure for many applications in cheminformatics including virtual screening ^7,8^. Molecular fingerprints, however, are computed from the chemical structure, e.g. as given in the form of SMILES ^9^ or InChI ^10^, which usually are only known for a tiny fraction of all mass spectra from complex mixtures ^11^. The most established approach to infer molecular fingerprints without known chemical structure is through support vector machines in combination with the computation of fragmentation trees ^12^, but this is computationally expensive, in particular for larger compounds. Recently, first attempts have been made to also use deep neural networks for directly predicting molecular fingerprints from mass spectra ^13,14^. One of the main obstacles in predicting molecular fingerprints is that they are typically large, very sparse, binary vectors. To further complicate things, the frequency of a fingerprint bit to be “activated” shows very large variations and makes it hard to learn how to correctly predict less common structural features (i.e., the bits). As a result, predictions only gave promising results for frequently activated bits of the molecular fingerprints^13^ and needed to be supplemented with closest matching library fingerprints to improve performance ^14^. With current open spectral libraries growing to such sizes that machine learning approaches have sufficient data for training, validation, and testing; there is an opportunity for the development of alternative mass spectral similarity scores.

Here, we present MS2DeepScore, a deep learning approach that is trained to predict structural similarities (Tanimoto or Dice scores based on molecular fingerprints) directly from pairs of MS/MS spectra without first computing molecular fingerprints. This is similar in spirit to the approach by Ji et al. ^15^ but uses a conceptually simpler Siamese neural network architecture ^16^. Furthermore, our approach only relies on peak m/z positions and intensities without requiring further spectrum information, as for instance the mass and chemical formula which are both necessary input for DeepMASS ^15^. The difference in input to the model also makes it difficult to directly compare these approaches quantitatively. Our proposed approach makes it possible to use MS2DeepScore for predicting structural similarities between spectra of various origins and with varying metadata quality. The model was trained using a dataset of 109,734 MS/MS spectra, which was built through curating and cleaning spectra obtained from GNPS ^17^ (see Methods). In addition to the prediction of a structural similarity, MS2DeepScore can also make use of Monte-Carlo dropout ^18^ to assess the model uncertainty.

We demonstrate that MS2DeepScore can predict structural similarities with high reliability. When comparing commonly used molecular fingerprints we achieve a root mean squared error for predicted Tanimoto scores of about 0.15 when run without uncertainty restrictions, and down to 0.1 with stronger restrictions on model uncertainty. MS2DeepScore is very well suited to detect compounds of high structural similarity and furthermore can create mass spectral embeddings that can be used for additional spectral clustering. We hence expect MS2DeepScore to become a key asset in building future MS/MS analysis pipelines. Depending on the desired application, MS2DeepScores could also be combined with other mass spectral similarity metrics to make full use of their complementary aspects.

## Results

A large set of MS/MS spectra was retrieved from GNPS^17^ and subsequently curated and cleaned using matchms^19^ (see Methods). The resulting training data set contains smiles/InChI annotations for 109,734 spectra, which allowed us to create molecular fingerprints to quantify structural similarities of spectral pairs. The dataset contains 15,062 different molecules (disregarding stereoisomerism - as represented by InChIKeys unique in the first 14 characters).

We randomly took out 500 of the 15,062 InChIKeys to form a validation set and again 500 to form a test set (see also Methods). The validation set (3,597 spectra of 500 unique InChIKeys) is used to monitor the model training process and explore the key hyperparameters while the test set (3,601 spectra of 500 unique InChIKeys) is used for a final unbiased evaluation of our model. Drawing pairs of spectra from the training set (102,536 spectra of 14,062 unique molecules), we trained a Siamese neural network to predict Tanimoto scores as depicted in Figure 1.

**Figure 1.**
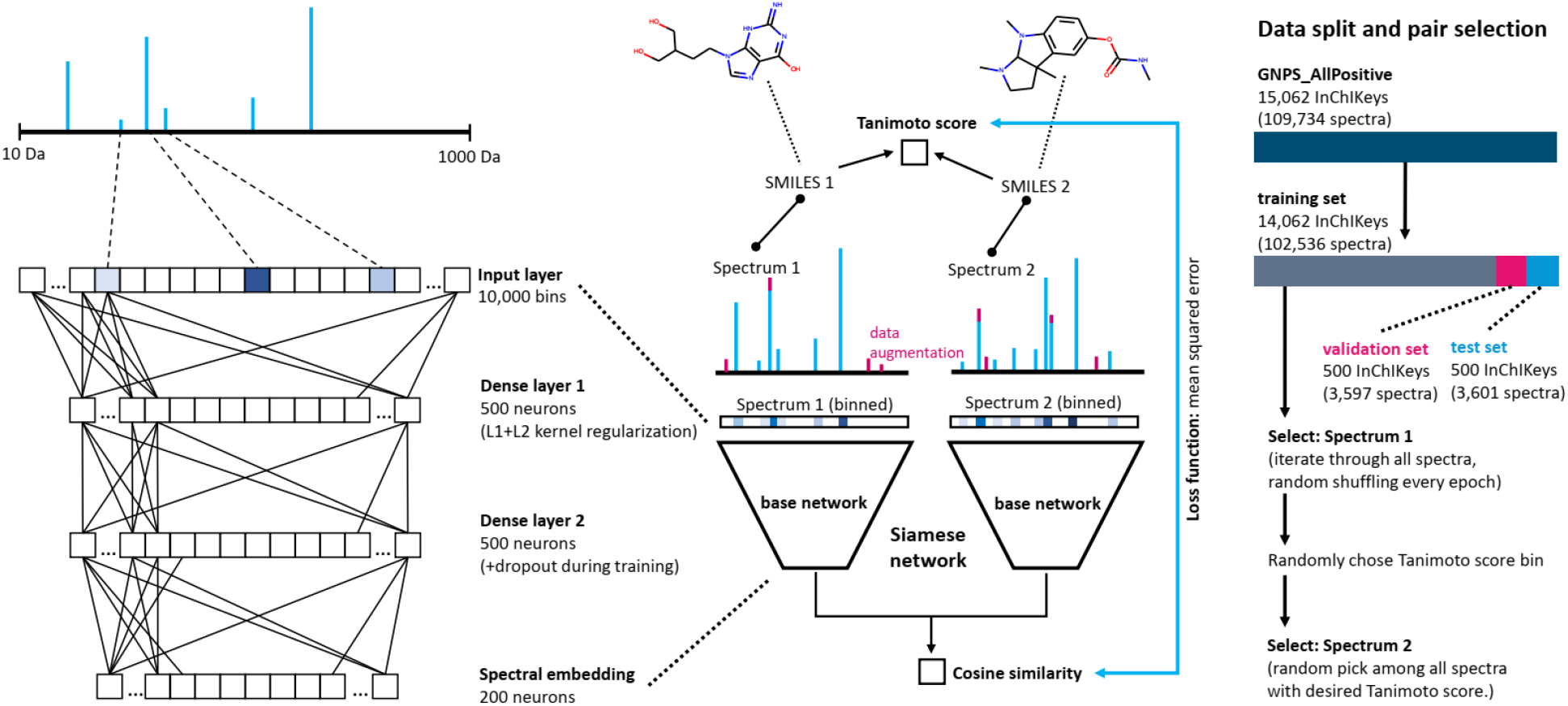
Sketch of the Siamese neural network architectures and training strategy behind MS2DeepScore. The Siamese network uses the same “base network” twice during training and prediction to convert a binned spectrum into a spectral embedding (200-dimensional vector). The network is trained on spectral pairs and the mean squared error between the cosine similarity of the two spectral embeddings and the actual structural similarity score (here: Tanimoto between rdkit 2048bit fingerprints). To increase the robustness of the model data augmentation techniques were used which includes moderate random changes of peak intensities as well as removal and addition of low intensity peaks.

A key challenge when training a neural network to predict Tanimoto scores is that the total set of possible spectrum pairs shows a highly unbalanced distribution in structural similarities, with most pairs displaying low structural similarity. Our procedure for drawing spectral pairs compensates for the unbalanced nature of the data, by selecting pairs with probabilities that are weighted according to their structural similarity, as described in detail in the Methods section. We further applied L1, L2 and dropout regularization, as well as data augmentation techniques to ensure the generalization of the model to unseen data.

### MS2DeepScore predicts Tanimoto scores with high accuracy

In real-world application of the model, it is acceptable that there is a small error in estimation of the structural similarity between two spectra, while outliers with large errors should be avoided. Therefore, the root mean squared error (RMSE) was used as an overall evaluation metric, since it penalizes large errors on individual samples. In addition, the model should ideally perform well across the full range of possible pair similarities, which for the here used Tanimoto and Dice scores lies between 0 and 1. However, the datasets are highly unbalanced in that respect, since most spectrum pairs have low Tanimoto scores (figure 3A). We hence decided to inspect the model accuracy not as a global average since that would strongly bias the outcome to the performance on low Tanimoto pairs. Instead, we split all possible spectral pairs into 10 equally spaced Tanimoto score bins. In figure 2A we display the distributions of the predicted Tanimoto scores for each bin, which reveals that the individual distributions show a high overlap with the correct Tanimoto scores. As expected, the prediction is not perfect. The distributions show long tails of predictions that differ from the true structural similarities. Looking at root mean squared errors (RMSE) across all Tanimoto score bins, it can be noted that MS2DeepScore generally performs very well and can predict Tanimoto scores between 0.1 and 0.9 with a RMSE between 0.13 and 0.2. Accuracy is lower for the highest and lowest Tanimoto scores, which may partly be attributed to the regression to the mean effect (the training loss makes it unattractive to approach the upper and lower score limit). The highest Tanimoto scores show a relatively long tail indicating more frequent wrong predictions (Fig. 2A). Predictions are also slightly more spread out for Tanimoto scores around 0.6-0.8, a range with relatively few occurrences and hence less training data in the dataset. Other underlying reasons can not be ruled out at this point, such as poorer correlation between fragmentation information and actual structural similarity scores. Taken together, MS2DeepScore is highly reliable in separating high, mid, and low structural similarity pairs (see also supplemental figure S1), but might be more error prone when it comes to smaller nuances.

**Figure 2.**
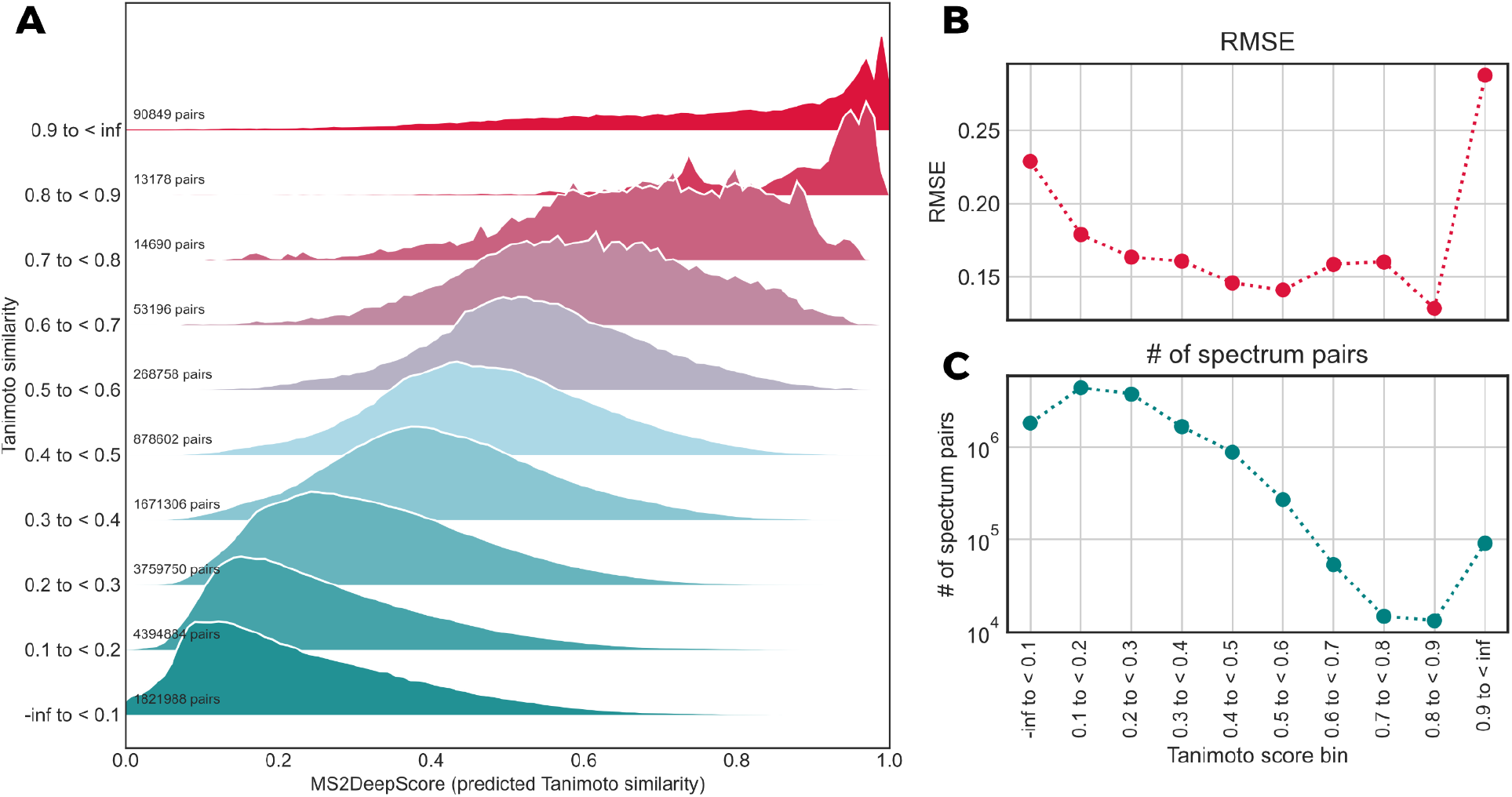
(**A**) To better account for the unbalanced nature of the total set of Tanimoto scores we here plotted the MS2DeepScore Tanimoto score predictions across different bins of Tanimoto scores (<0.1, 0.1-0.2 etc. until >0.9). (**B**) Average precision measures across the different Tanimoto score bins by RMSE. (**C**) The number of spectrum pairs which fall into each of the 10 Tanimoto score bins, illustrating the highly unbalanced nature of the dataset.

**Figure 3.**
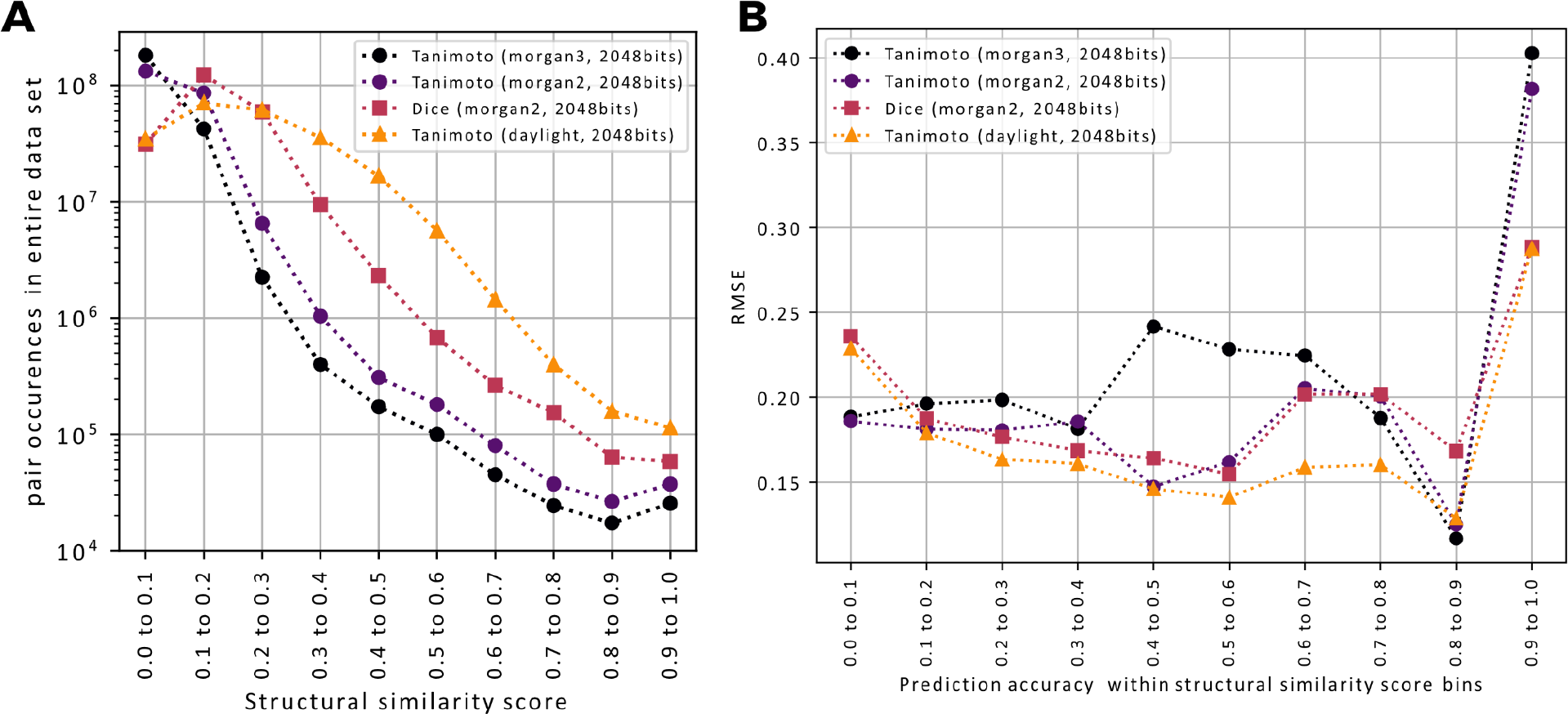
(**A**) Different molecular fingerprints (morgan2, morgan3, rdkit-daylight) and different scoring methods (Tanimoto/Jaccard vs. Dice) lead to very different distributions of scores across the full dataset (15,062 unique InChIKeys hence a total of 15,062^2^ pairs). Tanimoto scores based on rdkit-daylight fingerprints tend to give higher scores and thereby result in less unbalanced pair labels. The very pronounced unbalanced nature of Tanimoto scores on circular fingerprints (morgan2, morgan3) can partly be circumvented by switching to Dice scores instead. (**B**) MS2DeepScore models were trained for each different structural similarity score. RMSEs are here calculated for all spectrum pairs within the test set (3601 spectra) which fall into one of the 10 possible structural similarity score bins (x-axis labels).

### Accurate prediction of different structural similarity measures

Molecular fingerprints come in many different types and flavors which typically do not work equally well for all compound classes ^7,8,20^, and there is no general consensus on which molecular fingerprint to use (although MAP4 was recently said to “rule them all” ^20^) nor how to compute the actual similarity score from two fingerprints. Furthermore, even though Tanimoto (=Jaccard) is often considered as a standard metric ^21^, other metrics such as Dice are occasionally used. To show that our approach generalizes to a variety of structural similarity scores, we trained and tested the MS2DeepScore model for three different fingerprints (morgan2, morgan3 and daylight) as well as 2 different similarity metrics (Tanimoto and Dice). Overall, MS2DeepScore can make accurate predictions for all tested structural similarity measures (figure 3A). In addition, we observed that the precise distributions of all occurring structural similarity scores vary considerably when different structural similarity metrics are used (figure 3B). Structural similarity measures by Tanimoto scores from circular fingerprints (morgan-2 and 3, the rdkit^22^ pendants of ECFP-4 and ECFP-6) have a much higher tendency towards low scores when compared to Tanimoto scores from rdkit-daylight fingerprints. This can partly be adjusted by switching from Tanimoto to other metrics such as using a Dice score. Overall, we found that MS2DeepScore performs slightly better when predicting structural similarity scores with a less skewed distribution. This is to be expected since such scores display far more instances of moderate to high scores in the entire training dataset, for instance Tanimoto scores on daylight fingerprints result in 10-100 times more pairs across scores in the range from 0.2 to 0.9 when compared to Tanimoto scores on morgan-3 fingerprints (figure 3A).

### Detecting chemically related pairs: comparison to common mass spectral similarity metrics

Due to the low number of available correctly or reliably annotated mass spectra, many analysis pipelines have to rely on mass spectral similarity measures. A classical way to compare MS/MS mass spectra is to quantify the fraction of shared peaks as done by using variations of cosine similarity scores. Those measures tend to work well for very similar spectra, i.e. with many identical peaks. We recently introduced Spec2Vec, an unsupervised machine learning approach for computing spectrum similarities based on learned relationships between peaks across large training datasets ^6^. Spec2Vec based similarity scores were observed to correlate more strongly than classical cosine-like scores with structural similarities between the underlying compounds. An additional advantage is its fast computation, which allows to compare spectra against very large libraries. While trained on spectral data, Spec2Vec used an unsupervised method, meaning that it was trained on non-annotated data and did not make use of the structural information.

With MS2DeepScore, we now make use of the structural information that we have for a large fraction of the training data. Unlike Spec2Vec, which is trained to learn relationships between peaks from peak co-occurrences, and unlike modified Cosine, which computes the maximum overlap of matching peaks, MS2DeepScores is specifically trained to predict structural similarity scores. The ability of those different scores to identify chemical relatedness can thus not simply be compared by measuring their ability to predict Tanimoto score. In practice, however, all such scores are all used to identify chemically closely related compounds. Modified Cosine and Spec2Vec scores are for instance usd to generate molecular networks in GNPS ^4,6^. We therefore tested the scores ability to detect chemically related compounds by counting identified chemically related pairs within the test set (3601 spectra). Since “chemically related” is a hard to define concept, we simply operated with a fixed Tanimoto score threshold of 0.6 above which we call two compounds related. We then computed the precision and recall for finding structurally related compounds for all spectrum pairs above a threshold for the spectral similarity score which could be either MS2DeepScore, Spec2Vec, or modified Cosine (figure 4). This reveals that MS2DeepScore clearly outperforms both classical measures (two forms of the modified Cosine) as well as the unsupervised spectral similarity measure Spec2Vec, with respect to identifying high Tanimoto pairs, which can also be seen in the overall distribution of scores (figure S1). This makes MS2DeepScore a very promising approach for searching analogues in large datasets.

**Figure 4.**
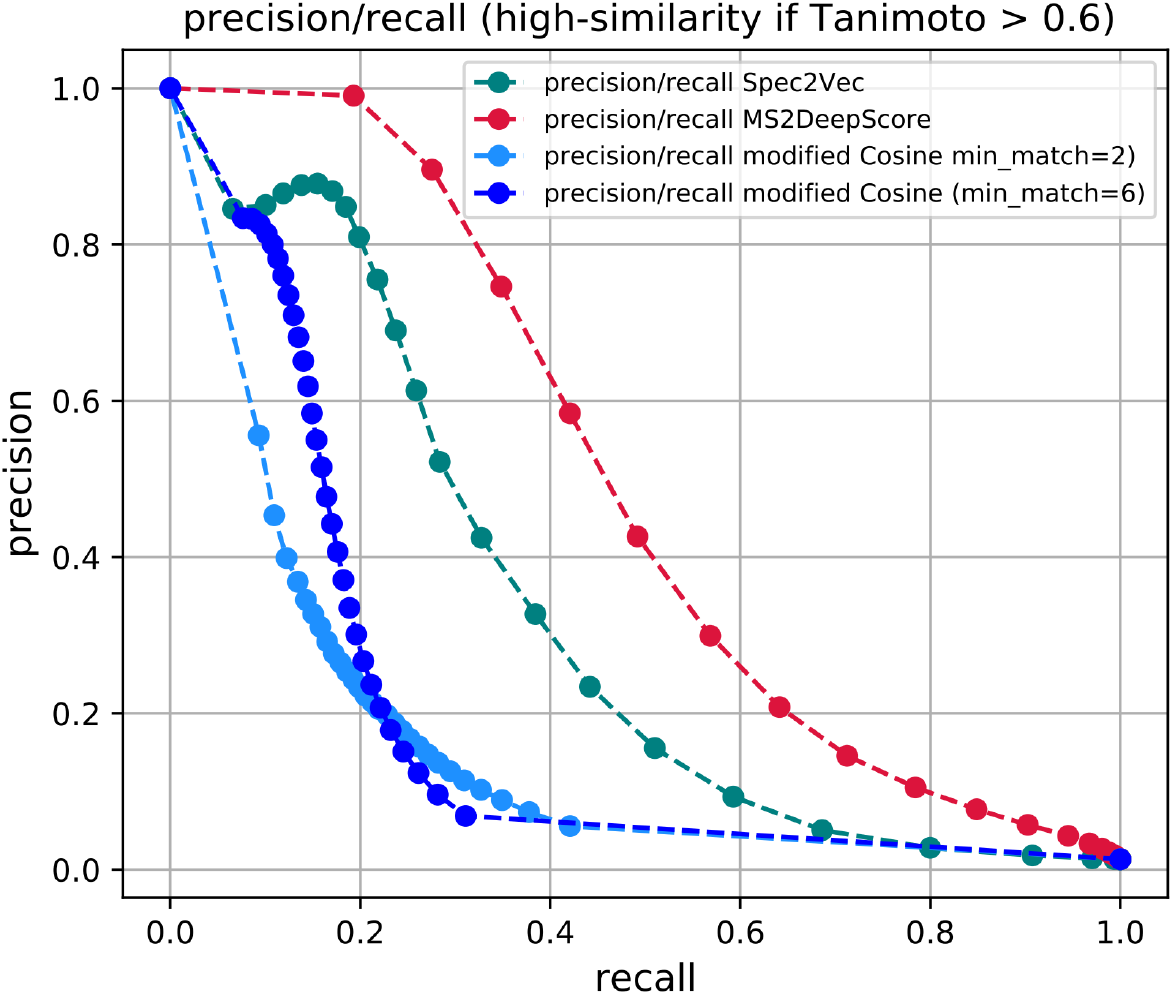
We here define a high structural similarity as Tanimoto > 0.6 and explore how well high structural similarity pairs can be retrieved using various spectral similarities. Collecting all spectrum pairs from the test set (3,601 spectra) with mass spectral similarity > X with X increasing from 0 to 1.0, we compute precision and recall for the different mass spectral similarity measures (MS2DeepScore, Spec2Vec, modified Cosine). The curves illustrate the tradeoff between higher recall (towards the right) and higher precision (towards the left). They also reveal that MS2DeepScore gives notably better precision/recall combination over the entire range, followed by Spec2Vec and only then modified Cosine.

### Combining different mass spectra for the same compound decreases Tanimoto score prediction error further

In many applications, such as library matching or analogue searching, datasets will frequently contain multiple mass spectra for a given compound. This is also the case for the data retrieved from GNPS (see Methods). The test set, for instance, contains 3601 spectra of 500 unique compounds (ignoring stereoisomerism). We hence tested whether structural similarity score predictions can be improved by taking the median of the scores calculated for different pairs of spectra corresponding to the same compound pairs. This can indeed be seen on the test set (figure 5, compare red and dark blue lines).

**Figure 5.**
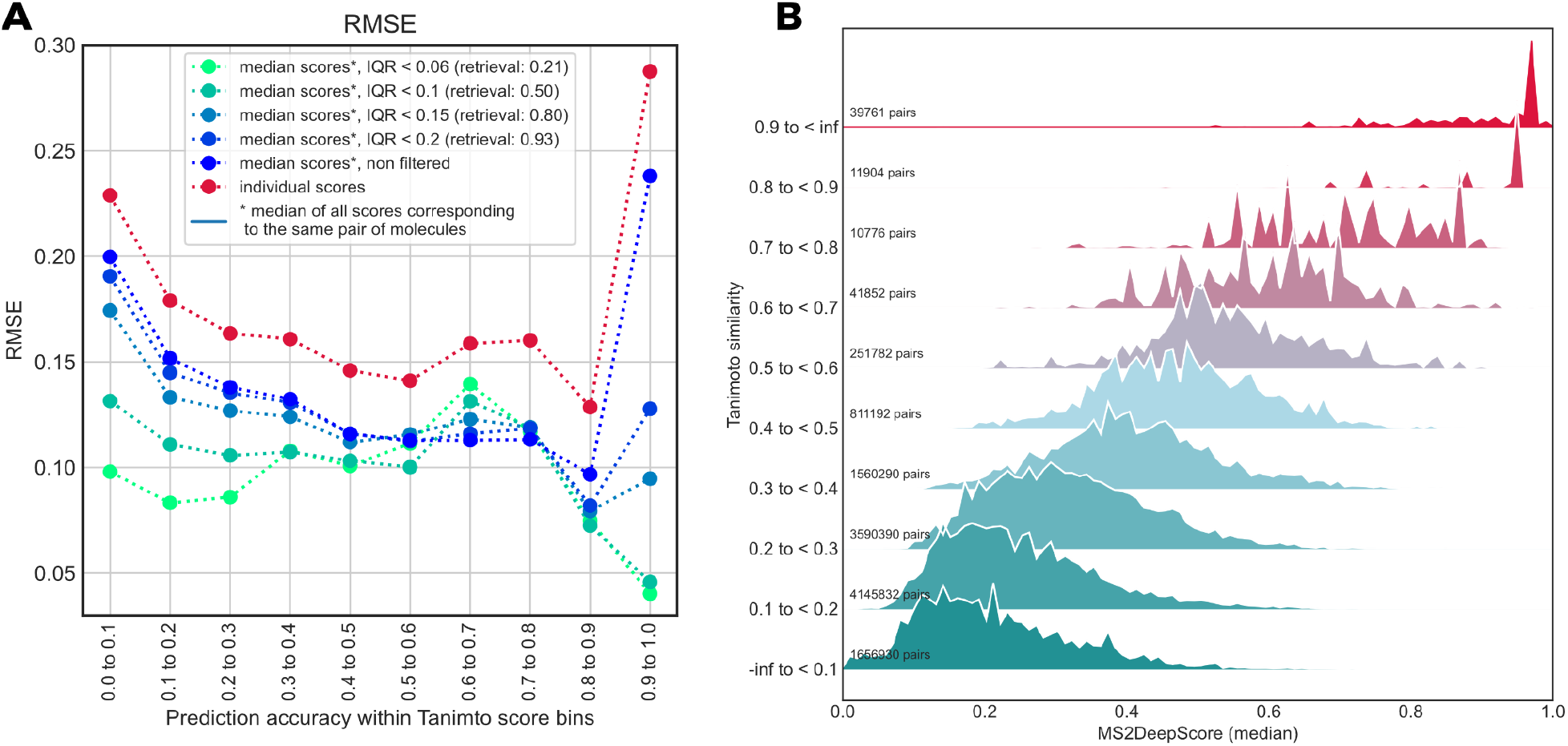
Combining MS2DeepScore predictions for spectral pairs corresponding to the same pair of molecules leads to more reliable Tanimoto score predictions. (**A**) Individual predictions(red dots) show consistently higher RMSEs than the median of predictions for (pairs of spectra corresponding to) the same compound pair (“all”). We also computed the interquartile range (IQR) of predictions of the same molecule pairs, which can be used to remove high IQR outliers. Retrieval rates after each label indicate the total fraction of scores that fulfilled the given criterium (IQR < x). (**B**) When compared to figure 2 it is apparent that the high Tanimoto score predictions become notably more reliable when removing scores with large variations of same-InChIKey predictions (here: keep scores with IQR < 0.2).

The improvement in accuracy becomes even more pronounced when removing potential outliers based on the interquartile range (IQR) of all predictions for the same pair of molecules. In particular high and low Tanimoto score predictions become notably more reliable, even at comparably high threshold IQRs (figure 5A, e.g. threshold < 0.2 which corresponds to 93% of all scores). Given the considerable improvement of the structural similarity prediction, we expect that this use of multiple predictions for mass spectra of the same compound pair can be applied successfully in practice, e.g. for library matching or analogue search, or when measuring multiple spectra of the same compound at various collision energies.

### Using Monte-Carlo dropout ensemble models to estimate prediction uncertainty

Using ensembles of multiple machine learning models is a frequently used technique for improving machine learning results, but also to assess the model uncertainty (also referred to as epistemic uncertainty). Ensembles can be built in many ways, but one particularly efficient ensemble learning technique for neural networks is Monte-Carlo dropout ^18^. It makes use of the dropout layers in a network to randomly silence a fraction of the nodes for each inference step. Traditionally, dropout is only activated during model training, but in Monte-Carlo dropout it stays in place when making actual predictions, which can be interpreted as a random sampling technique across a virtually unlimited set of model variations. Since the neural network architecture used for MS2DeepScore includes dropout layers (figure 1), it is straightforward to do such ensemble learning (figure 6).

**Figure 6.**
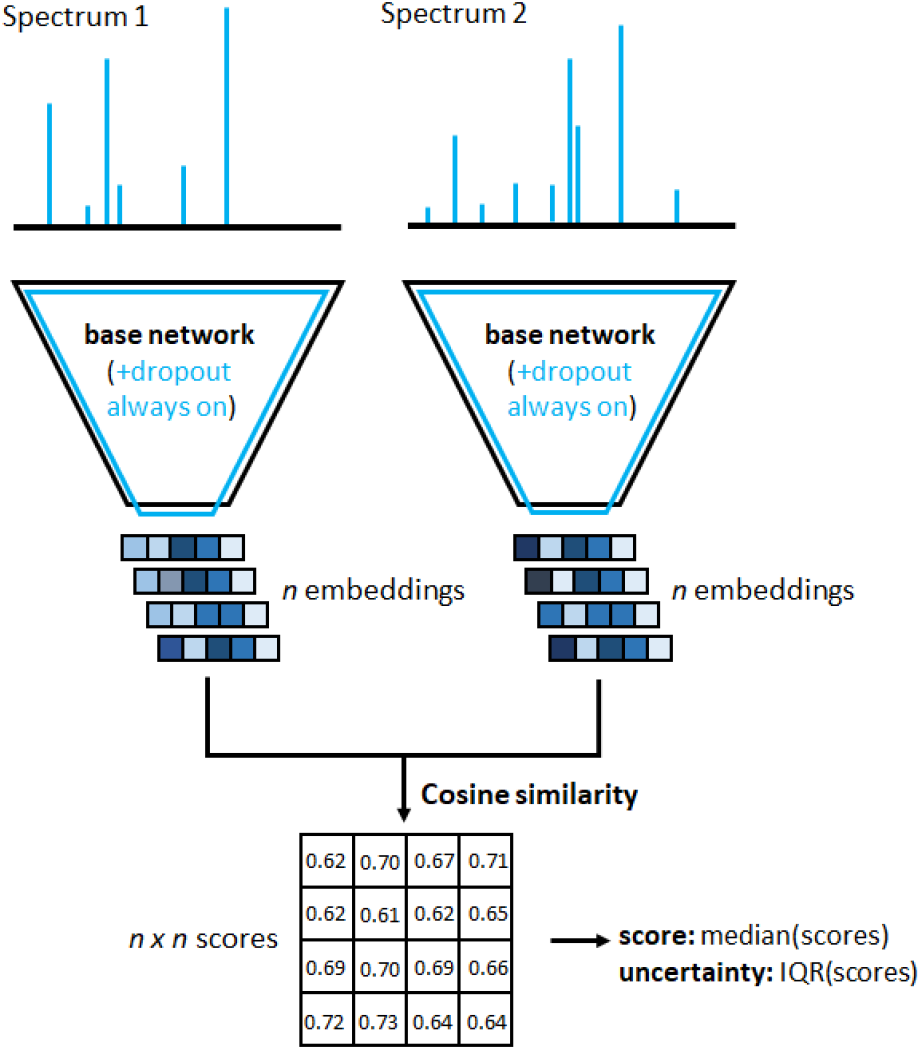
Sketch of MS2DeepScore running in Monte-Carlo dropout modus. By keeping the dropout layers switched on, model predictions will essentially be sampled from random variations of the respective neural network. With a dropout rate of 0.2, a random selection of 20% of all nodes in the central dense layer(s), see figure 1, will be silenced. From n resulting variations of the created spectrum embeddings an array of n*n scores can be computed. Finally, the array of ensemble scores is used to calculate a single median score together with the interquartile range as a measure of model uncertainty.

For a given spectrum, we compute *n* different embeddings, each from a slightly different version of the base neural network where 20% of its nodes are silenced (dropout rate=0.2). For a pair of spectra this results in n*n Tanimoto score predictions from which an ensemble score as well as a dispersion measure to assess the prediction certainty can be calculated. To be less sensitive to outliers, we chose to take the median score, rather than the mean score. The prediction uncertainty is measured by the interquartile range (25-75%) which is more suited than the median absolute deviation for non-symmetric distributions ^23^. This is also very accessible computationally since only *n* embeddings need to be generated per spectrum to obtain a total of n x n independent Tanimoto score predictions. Inference will hence only take 10x longer for an ensemble of 100 predictions.

We tested the resulting uncertainty estimate on all possible pairs within the 3,601 spectra of our test set. Taking the median of an ensemble of 100 scores already results in an overall drop in prediction error across nearly all Tanimoto score bins (figure 7, B, red vs. blue line). We then filtered out scores, according to increasingly stringent interquartile range (IQR) thresholds. Over the entire dataset, this approach leads to a large decrease in prediction error (figure 7, A) but comes at the cost of a lower retrieval. For instance, all predictions within IQR < 0.025 - which will discard about 75% of the scores - will result in a drop of the average RMSE from about 0.17 to about 0.11 (figure 7, A). It is important to note, though, that this average gain in precision is not distributed equally across the full range of Tanimoto scores. The RMSE drops most significantly in the low (<0.4) and high (>0.8) Tanimoto score range (figure 7, B), while the error slightly increases in the mid score range (0.5 - 0.7).

**Figure 7.**
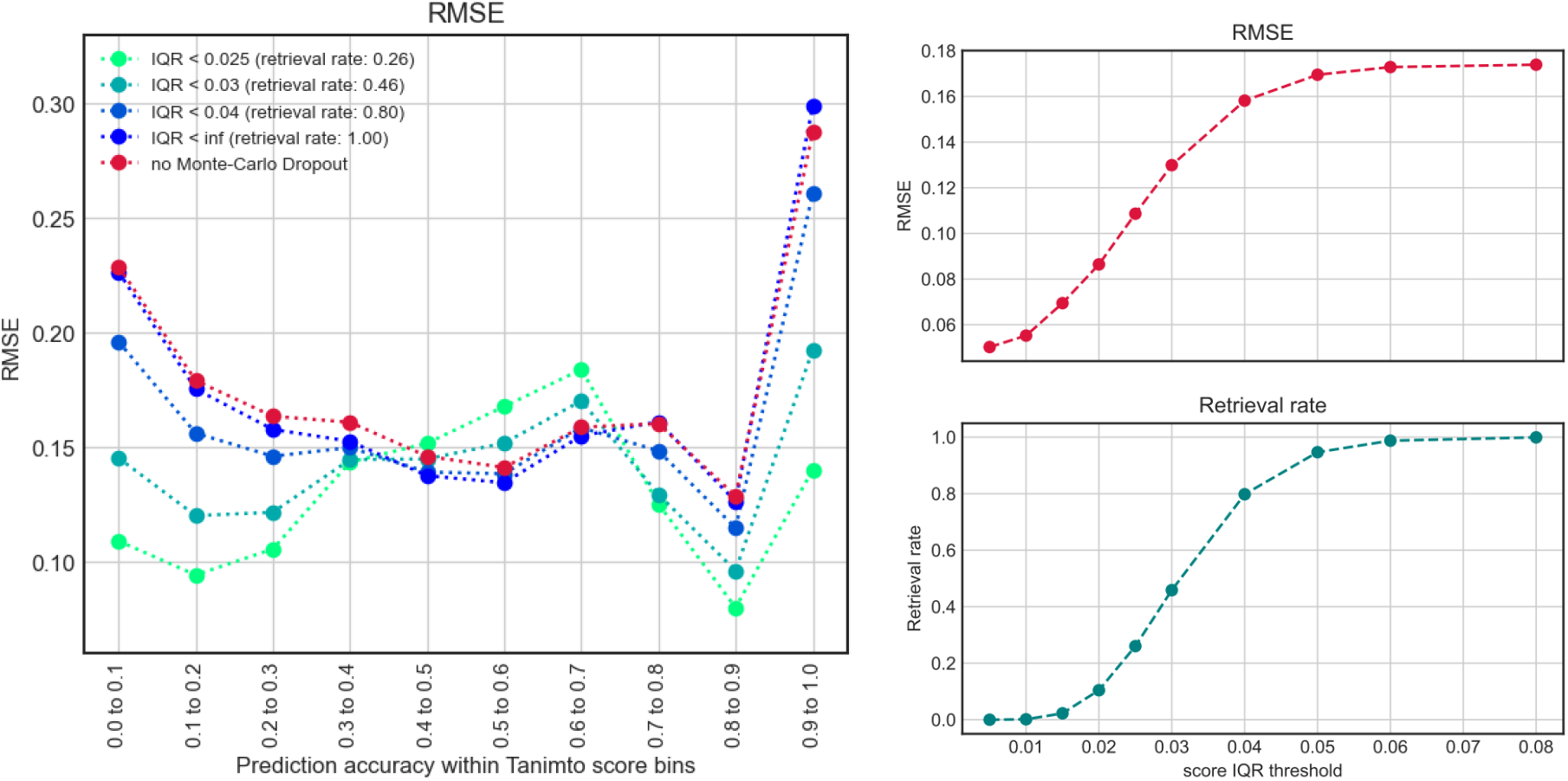
Monte-Carlo dropout provides Tanimoto score predictions, but also the interquartile range as an uncertainty measure (here over 10×10 individual scores). Discarding scores with higher uncertainties (higher IQR, interquartile range) does indeed improve the average prediction performance notably, although at the price of lowering the retrieval rate (retrieval rate = fraction of total scores with IQR < threshold, see lowest panel plot on the right).

### Embedding based mass spectral clustering

Unlike recent approaches to predict molecular fingerprints using deep learning ^13,14^, we have chosen to train a Siamese neural network^16^ to directly predict Tanimoto similarities. A key feature of our neural network design is the creation of abstract embedding vectors for each input spectrum (figure 1). This has two main benefits. First, it allows to scale similarity calculations much more efficiently to very large numbers of spectrum pair calculations by separating the mass spectrum embedding creation step from the actual similarity score calculation. The embedding creation includes the mass spectrum binning as well as the inference step with the ‘base’ neural network (figure 1) and is computationally far more expensive than the actual score calculation. Because embedding creation only needs to happen once for each spectrum instead of for each pair, this vastly reduces computational cost. As an example: predicting all possible similarity scores between the 3,601 spectra in the test set (6,485,401 unique pairs) took 5-10 minutes on a Intel i7-8550U CPU. The second reason for choosing this network architecture design is that such embeddings can have additional value beyond the Tanimoto score prediction. Even though they do not directly correspond to any conventional molecular fingerprint, they are trained to support a prediction of a fingerprint-based similarity score, and therefore we hypothesize that they will contain features that reflect chemical properties. This was tested by running t-SNE^24^ as implemented in scikit-learn^25^ on the 200-dimensional embeddings of the test set (3,601 spectra). This algorithm provides x,y-coordinates for every spectrum in the test set which we plotted and colored according to the 14 chemical superclasses provided by Classyfire ^26^ (figure 8a). Molecules of the same chemical class tend to cluster together in the resulting t-SNE plot, confirming that the MS2DeepScore embeddings represent molecular features. Figures 8b and 8c show that this conclusion also holds on a more detailed level, by zooming into a small region of the t-SNE plot (Fig 7b) and coloring according to Classyfire subclasses (Fig 8c).

**Figure 8.**
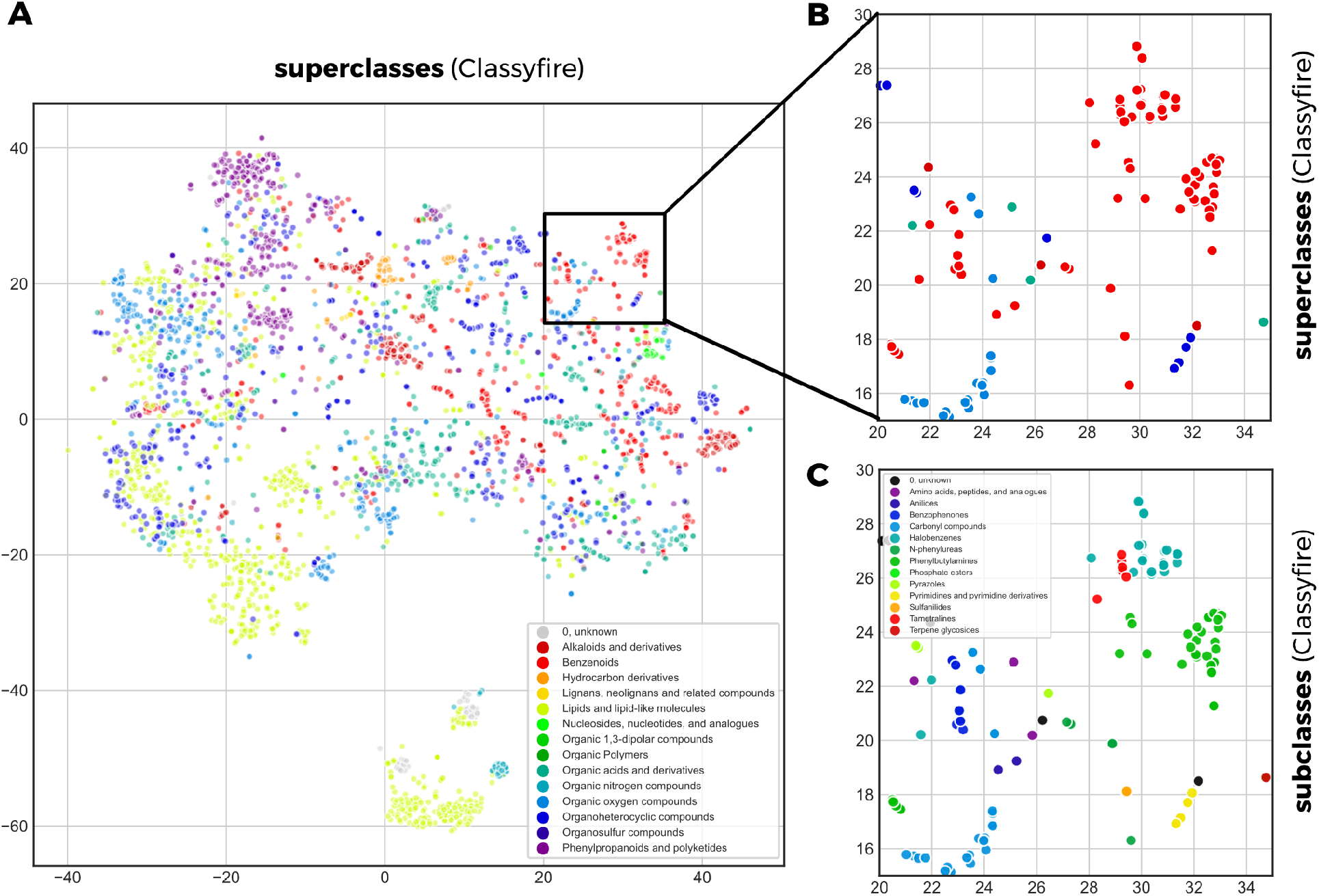
(**A**) 3601 spectra are plotted as colored dots with x,y positions derived through t-SNE based on the MS2DeepScore embeddings. Dots are colored according to 14 compound superclasses provided by Classyfire (large panel). (**B**) zooms into a small region for which (**C**) also the x Classyfire subclasses are displayed.

### MS2DeepScore Python library

MS2DeepScore is available as an easily installable Python library running on Python 3.7 and 3.8. Source code and installation instructions can be found on GitHub (https://github.com/matchms/ms2deepscore). The presented results were obtained using version 0.2.0. MS2DeepScore is integrated in matchms, a recently developed Python library for mass spectra import, handling and comparisons^19^.

## Discussion

Modern deep learning techniques have quickly gained popularity in many research fields and in some cases even started to replace classical, more heuristic techniques (e.g. in computer vision and natural language processing). The application of deep learning on fragmentation mass spectrometry data though, has only just begun to enter the stage. The first promising applications include the prediction of compound classes from MS/MS spectra ^27^ or from (predicted) molecular fingerprints ^28,29^, the prediction of bioactivity signatures^30^, the prediction of parts of molecular fingerprints ^13,14^, as well as the prediction of the structural similarity from MS/MS spectra and chemical formula ^15^. With MS2DeepScore, we show for the first time that neural networks can also be used to predict structural similarity scores, i.e. a chemical-driven measure, from MS/MS spectra without requiring a known molecular formula or other metadata. We found that predictions are generally accurate (MAE of about 0.12) and in particular get the general tendency right with large outliers being rare (RMSE of about 0.15). By constructing the test, validation and training sets from separate sets of molecules we show that the presented MS2DeepScore models are predictive for novel molecules. Selecting all spectra for 500 randomly chosen compounds for our test set should reflect the overall diversity of the MS/MS dataset well enough. In addition, we observed that the distribution of Tanimoto scores within the test set shows a similar profile as the distribution for the training set (figure 2B vs. figure 3A) and that the chemical diversity in the test set is high, with 14 chemical superclasses, 99 different chemical classes and 140 different chemical subclasses found via Classyfire ^26^.

MS2DeepScore comes with two inherent downsides, when compared to conceptually simple, heuristic measures such as the Cosine spectral similarity score. One limitation that is common for all machine learning-based approaches is that a score itself is not deterministic, but might change when a new model is trained (e.g. when using different parameters or different training data). In practice that can often largely be addressed by using properly versioned, pre-trained models. The second downside that is typical for neural networks is the lack of an intuitive explanation of *why* a certain score is given, a common problem in the field of deep learning (i.e., explainability or explainable AI). Here, however, we also provide the option to use Monte-Carlo Dropout, an ensemble learning technique which makes it possible to assess the model’s own uncertainty. This should in practice help in better identifying and handling uncertain predictions, e.g. for data that are very different from the training data.

Another possible limitation of MS2DeepScore comes from the maximum achievable precision. Even after numerous experiments with training “deeper” neural networks (with more layers), wider neural networks (more nodes per layer), or less restrictive mass binning (more m/z bins), we could not significantly decrease the overall Tanimoto prediction error (supplemental material). Obviously, that does not prove that there cannot be a better performing neural network for the given task. However, it strongly indicates that the achieved precision is already fairly high, given the many limitations of the used dataset. These limitations arise from the fact that the used spectra come from different instrumentation types, are of varying quality, and are likely to be very unbalanced with regard to represented compound classes and types. It was, however, possible to reduce prediction errors considerably by applying various ensemble learning techniques. Applying Monte-Carlo Dropout (figure 7) or using ensembles of different architecture models (supplemental, figure S2) lead to more reliable results and provide means to assess the prediction uncertainty which allow users to further specify the desired level of precision. The same was seen when combining scores for spectra obtained for the same pair of molecules(figure 5).

It is important to note that the neural network was not trained on any spectrum metadata such as parent mass and elemental formula, like for DeepMASS^15^. Such metadata could include parent mass, precursor ion charge, adduct information, instrumentation type, spectral quality, or more processed information such as the elemental formula or chemical compound class. We speculate that incorporating relevant metadata in the pairwise predictions would have increased the accuracy in our evaluation. However, here, we chose not to include such metadata. Having a way to predict structural similarities solely based on MS/MS peaks allows MS2DeepScore to be easily applied to a large number of spectra without costly and timely spectral processing or matching steps. If users have access to this metadata, they can anyway still use it for an independent selection step or to train an additional small model for removing likely outliers based on metadata pairs. Furthermore, the use of just mass fragments also makes MS2DeepScore applicable to GCMS data which generally lack precursor masses.

We expect that another promising route to further improve the predicted scores lies in using complementary aspects of different spectral similarity scores. MS2DeepScore usually comes very close with its Tanimoto score predictions, but might not always be precise enough to handle all nuances. It is - for instance - difficult to discriminate between high Tanimoto scores (say 0.8-0.9) and a near-complete chemical match. Reliable identification of exact compound matches hence requires additional algorithms, such as the successful use of machine learning in combination with computational fragmentation trees combined with library data^31^. Key advantages of MS2DeepScore over such an approach are the very large gains in computation time which will allow to run very extensive screenings between many thousands of compounds, and its ability to predict structural similarities based on spectra of novel molecules without having to use any metadata or having to consult with library data. In practice, we expect hybrid approaches that combine multiple algorithms to be a promising route forward. MS2DeepScore could be applied for preselecting candidates prior to a computationally more expensive step (e.g. using fragmentation trees) or it could be combined with other similarity scores to improve the prediction reliability, e.g. by also consulting other scores such as cosine-based spectral similarity scores or Spec2Vec. The latter is frequently outperformed by MS2DeepScore (figure 5 and S1) but - as an unsupervised approach - has the advantage that it can be trained on unlabeled data which is not directly accessible for the presented supervised approach.

Being able to predict Tanimoto scores, or more precisely Tanimoto scores computed from Daylight2048bit fingerprints available in RDKit, can be interpreted as being able to infer chemical relatedness from MS/MS spectra. There is, however, no consensus on how to best quantify chemical relatedness which resulted in a large variety of different molecular fingerprints^7,8,20^ as well as fingerprint based scores^21^. We showed that MS2DeepScore can be trained to predict various different scores such as Tanimoto on Daylight fingerprints, Tanimoto on morgan-2 (similar to ECFP-4), or Dice scores on morgan-2. Given the huge variety of fingerprint types, their dimension (number of bits) as well as the used scoring metrics, our explorations should only be seen as a starting point, but our observations already suggest that MS2DeepScore will be able to cope reasonably well with a large variety of different fingerprints and metrics (figure 3). Since our neural network creates low dimensional embeddings for each spectrum it will also be possible to combine different structural similarity measures by stacking embeddings of different models that were trained on different scores. Another future path might be to modify the Siamese network architecture to predict various structural similarity scores at the same time. To fuel future improvements on structural similarity prediction of MS/MS spectrum pairs it is vital that the field converges onto a standardized way of evaluating and comparing approaches.

MS2DeepScore comes as an easy to install and easy to use Python library and the actual scores are fast to compute. In particular the ability to split spectral embedding creation from similarity score calculation makes it very scalable to large-scale comparisons (many thousands of spectra). Our MS2DeepScore model which was trained on a public dataset of about 100,000 spectra of 15,000 compounds can be found online (see link in Methods). Even though we here showed that such a model performed well on spectral data of unseen compounds, it is to be expected that training on even larger, more diverse, or more curated datasets will further improve the model performance. In this light, training (and ideally: providing) new MS2DeepScore models on alternative spectral libraries such as METLIN^32^ or NIST^33^ could become an important step to improve neural network based predictions.

Finally, we speculate that MS2DeepScore-generated spectral embeddings can further be used for other fascinating tasks. Here, we illustrated its ability to position spectra into chemically meaningful clusters (figure 8). This also makes MS2DeepScore a promising complementary candidate to the mass spectral similarity metrics used in established mass spectrometry based network analysis and clustering tools such as GNPS ^17^ or MetGem^34^.

## Conclusions

MS2DeepScore is a deep learning technique to predict structural similarity scores between fragmentation mass spectral pairs. We show that MS2DeepScore is able to infer structural similarities between mass spectra with high overall precision, without requiring any additional metadata or library data. We demonstrate that the accuracy of the predictions can be improved notably by the use of various ensemble learning techniques, in particular by merging predicted scores of spectra belonging to the same compound pair or by applying Monte-Carlo Dropout to sample from random model variations.

MS2DeepScore is very fast and scalable. We conclude that this makes MS2DeepScore a powerful novel tool for running large scale comparisons and analyses, for instance on complex mixtures rich in spectra of unknown compounds. We expect that MS2DeepScore can generally be used to complement -or replace-common currently used spectral similarity measures in many metabolomic workflows, including network analysis and clustering approaches.

## Methods

### Data and data preparation

#### Spectrum data preparation

We use LCMS (MS/MS) spectra from GNPS, which underwent basic metadata cleaning as described in ^6,19^. The dataset was retrieved from GNPS (25/01/2021) and contains a total of 210,407 MS/MS spectra. Metadata was cleaned and checked using matchms^19^ version 0.8.2, which included cleaning compound names, extracting adduct information from the given metadata, moving metadata to consistent fields and conversions between InChI and SMILES as well as to InChIKeys when missing and when possible. We then ran an automated search against PubChem^35^ using pubchempy^36^ for spectra which still missed InChI or SMILES annotations. The full cleaned dataset (210,407 spectra, 184,698 annotated with InChIKey and SMILES and/or InChI) can be found on zenodo: https://zenodo.org/record/4699300.

We here focus on spectra acquired in positive ionisation mode with proper InChIKey as well as a SMILES and/or InChI annotation, which in addition must contain >= 5 peaks between 10.0 and 1000.0 Da. This resulted in 109,734 spectra with 15,062 unique InChIKeys (considering only the first 14 characters).

The spectra underwent basic filtering to remove excessive amounts of peaks, by removing peaks with intensities < 0.1% of the maximum peak intensity and limiting the maximum number of peaks to the 1000 highest intensity peaks. This is mostly done to speed up the later binning and training steps and the hence removed peaks are most likely noise peaks. Peak intensities were square root transformed to avoid a too strong focus on the highest intensity peaks only. Spectrum peaks were binned in 10,000 equally-sized bins ranging from 10 m/z to 1000 m/z. In case multiple peaks ended up in one bin the highest peak intensity was chosen as value for that bin. Bins that were not filled in any of the training-data spectrum representations were removed from the vector representation, here this meant that 9,948 out of 10,000 possible bins are known to the model. The resulting vector-representation of the spectra served as input for the model.

#### Structural similarity label for spectrum pairs

Unless noted otherwise, we used Tanimoto scores on rdkit^22^ daylight fingerprints (2048bits) to compute structural similarities. For every unique 14-character InChIKey the most common InChI was selected (if different InChI existed) and used to generate a molecular fingerprint (as implemented in matchms^19^). For each pair of molecular fingerprints Tanimoto scores were calculated, indicating the structural similarity of that pair. This resulted in a matrix of 15,062 x 15,062 Tanimoto scores to be used as labels for the model training.

**Figure S1 with: 1. distribution of intensities before and after transformation. 2. Distribution of Tanimoto scores in the dataset before and after balanced data generation.**

#### Data generation

The set of 15,062 InChIKeys was split into a training (n=14,062), validation (n=500), and test set (n=500). To feed the data to the model effectively it was key to solve 2 challenges: 1) The structural similarity label distribution for all pairs is heavily left-skewed (most pairs are not similar, see figure 3A). 2) Per unique InChIKey multiple spectra could be used. Our MS2DeepScore Python library offers two types of data generators, one which iterates over all unique InChIKeys (*DataGeneratorAllInchikeys*) and one which iterates over all spectra and was used for the presented work (*DataGeneratorAllSpectrums*). The following algorithm was used to generate one cycle of training data, in each training epoch we used one cycles (i.e. we went through all spectrums in the training set once): Each spectrum was then matched to a random other spectrum, with the condition that the resulting corresponding InChiKey pair had a structural similarity label falling into a randomly chosen bin, which in our case were 10 equally-sized bins between 0 and 1. In cases where the structural similarity label for none of the pairs fell into the selected bin, the bin was iteratively widened by 0.1 until a structural similarity label fell into the bin.

After every training epoch, the loss on the validation set was computed. As for the training data we here used DataGeneratorAllSpectrums on the validation set. To ensure dataset consistency across experiments we used a fixed random seed for the validation set. We also used 10 cycles for the validation set which means iterating 10 times over all 3,597 spectra to monitor the training progress on a total of n=35,970 spectrum pairs. For the final evaluation on the reserved test set, we used all possible spectrum pairs between the 3,601 for the test set (n= 6,485,401 unique spectrum pairs).

#### Data augmentation

To ensure the model generalizes well to the test dataset and avoid overfitting we applied three forms of data augmentation on the binned spectra. 1) **low-intensity peak removal**: For a randomly chosen percentage (in the range of 0-20%) of non-zero bins with an intensity below 0.4 (actual intensity before transformation) the intensity was set to 0. 2) **peak intensity jitter:** Each non-zero bin intensity (after transformation) underwent changes between 0 and +-40%. 3) **new peak addition:** For each of between 0-10 randomly selected zero-intensity bins that bin’s intensity was set to random values between 0 and 0.01 (after transformation). Data augmentation was applied for every training example during training data generation.

### Deep learning implementation

#### Network architecture

We train a deep learning network on pairs of MS/MS spectra to predict the respective structural similarity label. For this, a Siamese network is used ^16^ which has 2 components: 1) A base network that creates abstract embeddings from both input spectra, and 2) A “head” part of the Siamese network which consists of a cosine calculation between both embeddings (figure 1). In the base network, the binned spectrum vector is passed through a series of densely connected layers until an abstract embedding vector of desired dimension is created as output. Based on a screening of various key parameters (see supplemental information) we settled on an architecture as depicted in figure 1: binning spectrum peaks between 10.0 and 1000.0 Da into maximum 10,000 same-width bins. This input vector is then followed by 2 densely connected layers, each with 500 nodes, followed by a final dense layer of 200 nodes for creating the spectral embedding. Two key measures are taken to prevent overfitting and improve generalization of the model to unknown data. Firstly, modern regularization techniques are applied ^37^. The deep neural network is trained using L1 (10^-6^)and L2 (10^-6^) weight regularization in the first dense layer, as well as dropout in the subsequent layers (dropout rate=0.2). In addition, batch normalization is applied after each dense layer except the output layer.

#### Uncertainty quantification using Monte-Carlo Dropout ensembles

To estimate the uncertainty of a prediction we used Monte-Carlo Dropout ensembles ^18^. At inference time, dropout was applied to all but the first layer of the base network. N=10 embeddings were created from an ensemble of these networks with dropout enabled. This resulted in a distribution of structural similarity predictions, for which the median and interquartile range (IQR) were calculated.

#### Training details

Models were trained with the Adam optimizer ^37,38^ that optimized the mean squared error (MSE) loss. We used a batch size of 32, and a learning rate of 0.001. Training continued until the validation loss did not decrease for 5 epochs (early stopping). Model training was done on GPU nodes from SURFsara with nvidia GTX 1080 Ti graphic cards (Lisa cluster). The fully trained model used to create figure 2, 4, 5, 7, 8 can be downloaded from zenodo: https://zenodo.org/record/4699356

#### Precision/recall analysis for selecting high Tanimoto score pairs

The precision/recall plot in figure 4 was created by measuring how many pairs with Tanimoto scores above a set threshold (=”high structural similarity pair”) were among a subset of all pairs for which the spectral similarity score was >*threshold_score*. We varied the threshold_score from 0 to close to 1 and recorded the precision and recall. By precision we here understand the number of high structural similarity pairs in the selection divided by the number of all selected pairs. Recall refers to the number of high structural similarity pairs in the selection divided by all high structural similarity pairs.

#### T-SNE on mass spectral embeddings from MS2DeepScore

For figure 8, we used the MS2DeepScore base network (figure 1) to compute the 200-dimensional spectral embeddings for all 3,601 spectra in the test set. Using the t-SNE^24^ implementation from scikit-learn^25^ we computed two-dimensional coordinates for all spectra. Here we used the following settings: metric=‘cosine’, perplexity=100, learning-rate=200 (default) and 1000 iterations (default).

## Acknowledgements

This work was carried out on the Dutch national e-infrastructure with the support of SURF Cooperative. J.J.J.v.d.H. acknowledges funding from an ASDI eScience grant, ASDI.2017.030, from the Netherlands eScience Center.

The authors thank the GNPS community for contributing annotated spectra to the public GNPS spectral library.

