## Supplemental materials for "MS2DeepScore - a novel deep learning similarity measure for mass fragmentation spectrum comparisons"

### Score comparisons with respect to precision-recall in highly unbalanced dataset

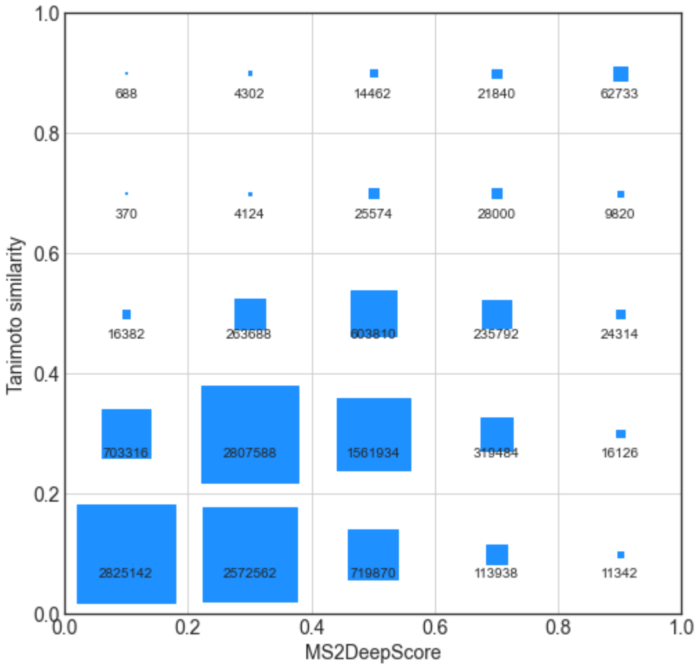


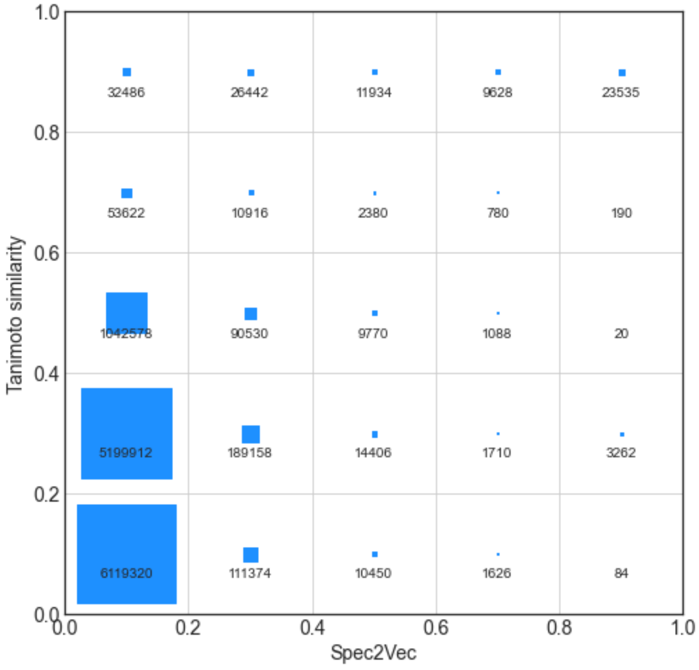

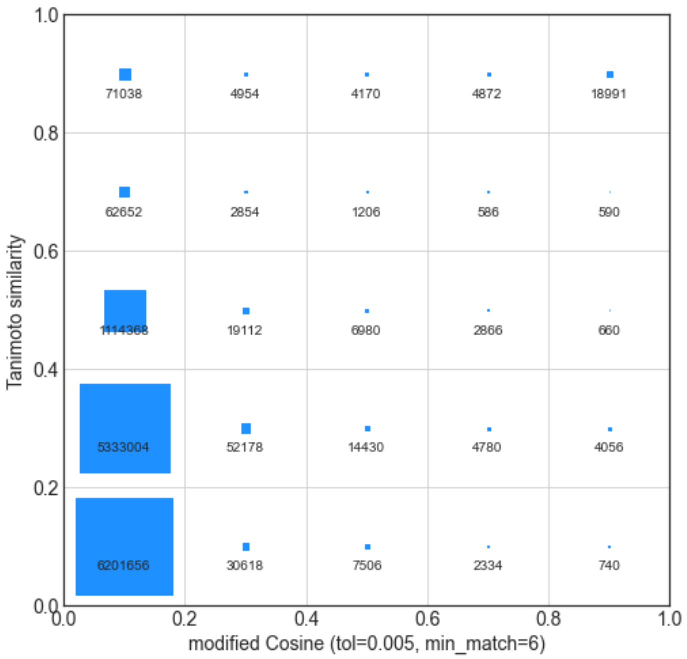


**Figure S1**. Similar to a confusion matrix for evaluating classification tasks, this plot displays the total fraction of all scores (3601 x 3601 possible pairs) which falls into one of 5 Tanimoto scores bins and one of 5 predicted Tanimoto score bins. The area of the squares reflects the total amount which is also given as absolute numbers.

### Hyperparameter search

#### Spectrum binning

To be able to handle the sparse MS/MS spectra, we only keep spectrum peaks within m/z values between 10.0 and 1000.0 Da and apply a binning step to reduce the possible number of m/z positions to a desired value. There is an obvious trade-off between more peaks which better reflect peak m/z positions and less bins which drastically reduces the necessary network parameters and computational performance. We were, however, surprised to observe a rather small impact of the spectrum binning on the overall Tanimoto predictions as measured using the root mean squared error (RMSE, Figure S1, top) or the mean absolute error (MAE, Figure S1, bottom). Here, we only varied the input dimension from 500 to 15,000 bins and kept the other network training parameters constant.
The fact that the initial binning has a rather moderate influence on the prediction accuracies gives a first hint at what information is key for the neural network to make a Tanimoto score prediction. Apparently, the larger pattern of the main peaks (i.e. relative positions and intensities) is far more important than

the precise m/z locations. For very low numbers of bins (<1000)  the prediction accuracy worsens notably and the predicted Tanimoto scores can hardly discriminate nuances in the range between 0.6 and 1.0 (see figure S4). Accuracy keeps improving when binning is becoming finer, but naturally this also results in a rapidly increasing number of model parameters, we decided to keep the input dimension at 10,000 bins.

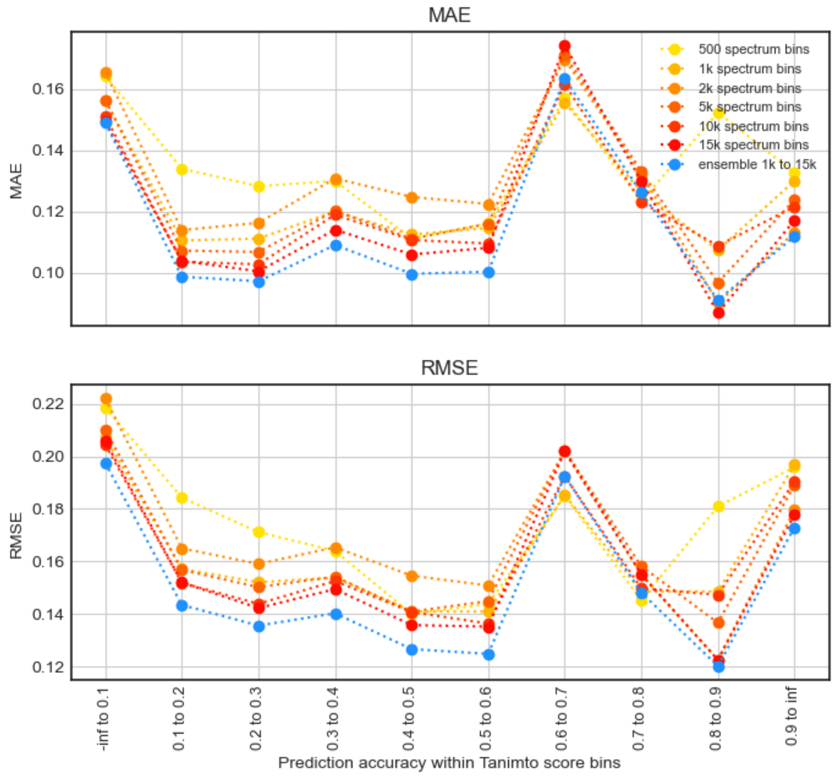


**Figure S2.** Different MS2DeepScore models were trained for different spectrum compression ranging from 500 to 15,000 bins (considering peaks in the m/z range from 10.0 to 1000.0 Da). In addition, an naive ensemble model was tested which takes the median prediction of the 5 models between 1k and 15k bins, blue dots). The MAEs and RMSEs are here calculated for all spectrum pairs within the validation set (3597 spectra) which fall into one of the 10 possible Tanimoto score bins (x-axis labels).

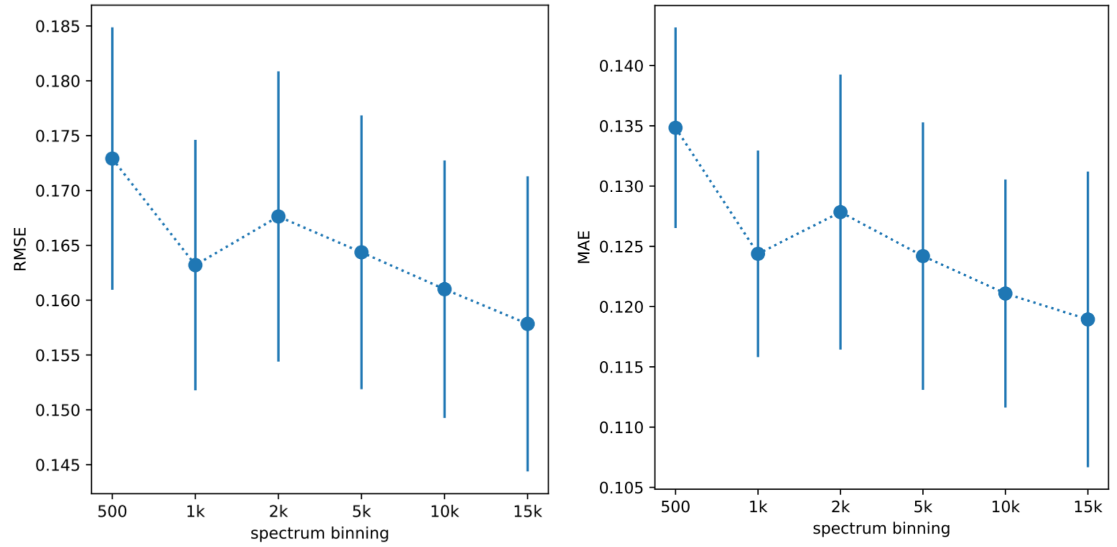


**Figure S3.** Spectrum were binned to values between 500 to 15,000 bins while all other network training parameters were kept constant. Models were training using early stopping with a patience of 5 epochs. Scores were computed for all possible pairs of spectra within the validation set and compared to the actual Tanimoto scores within 10 bins from 0 to 1.0 to account for the unbalanced nature of the dataset (see figure S2). The respective mean values across the 10 bins are plotted here together with the standard deviation (error bars are -0.5 STD to +0.5 STD). (Left plot) Mean RMSEs for different spectrum binning. (Right plot) Mean MAEs for different binning.

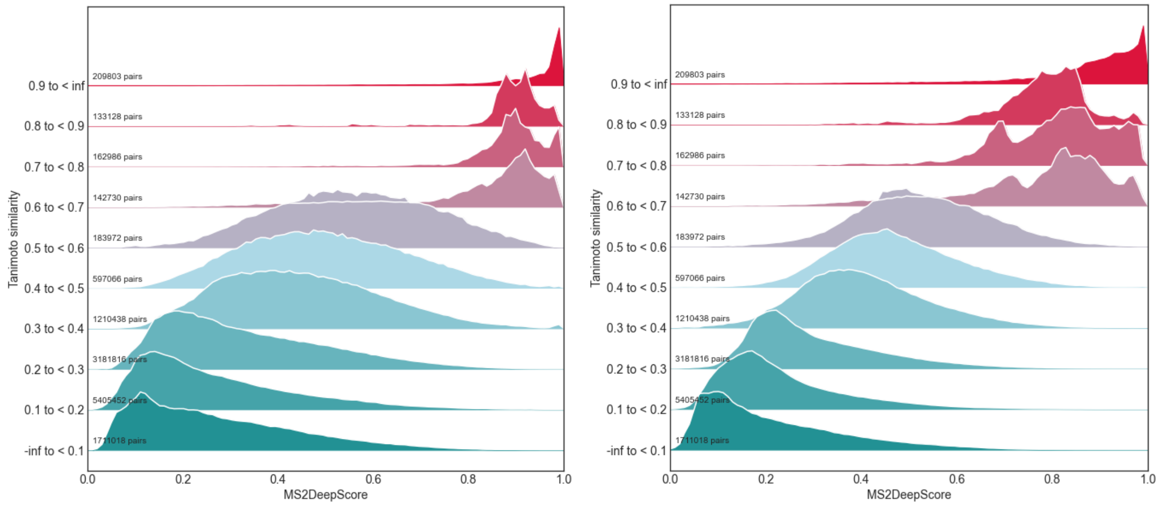


**Figure S4.** Comparisons between predicted Tanimoto scores (MS2DeepScore) and Tanimoto score (rdkit daylight fingerprint, 2048bits), for 500 spectrum bins (left) and 15,000 spectrum bins (right).

#### Network depth

Different numbers of densely connected layers were tested. The base network contains a first compression step from dim_bins → dim_01 followed by a number of densely connected layers, and finally an embedding creation → dim_embedding. We tested architectures ranging from 2 layers (10k→ 500→ 200) to 5 layers (10k→ 500 → 500 → 500 → 500 → 200) without achieving any notable improvement when using more than three layers (figures S5 and S6).

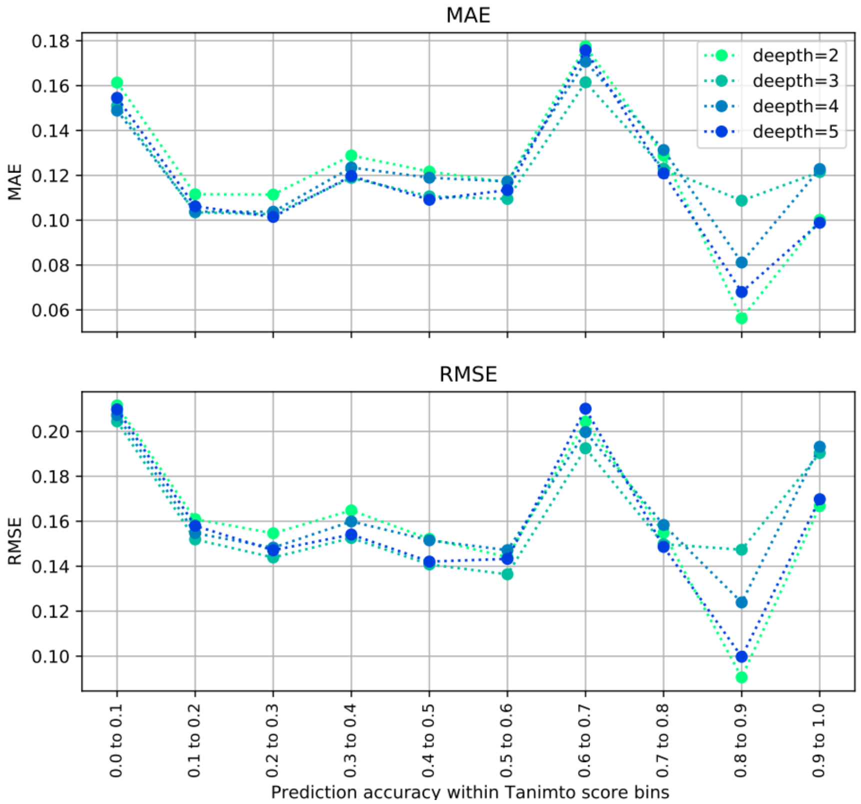


**Figure S5.** Different MS2DeepScore models were trained for different network depths, ranging from 2 to 5 dense layers. The MAEs (upper plot) and RMSEs (lower plot) are here calculated for all spectrum pairs within the validation set (3597 spectra) which fall into one of the 10 possible Tanimoto score bins (x-axis labels).

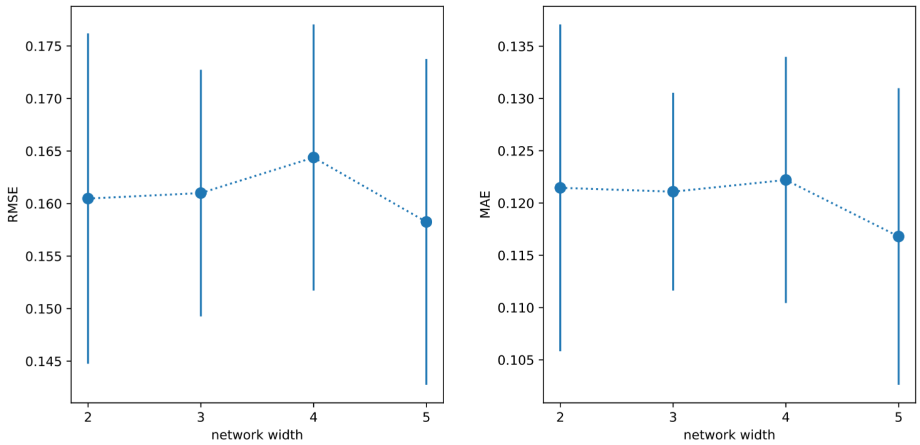


**Figure S6.** Plot showing the mean values for the RMSEs (left) and MAEs (right) for MS2DeepScore models with 2 to 5 dense layers. Means are simply the means across all 10 Tanimoto score bins shown in figure (figure S5). Error bars range from -0.5STD to +0.5STD based on the RMSE for all 10 Tanimoto bins.

#### Network width

Analogue to the network depth we also tested different network widths, i.e. different number of nodes per dense layer. Mostly due to the large number of weights for the first compression step, the total number of parameters is increasing rapidly for wider networks.

The prediction error generally increases for less nodes, but only really breaks down for very small networks (figure S7, e.g. 10k→ 20→ 20→ 200). It also seems to saturate for a width of about >1000 nodes. To us, this suggests that the key information is extracted in the first network layer where the sparse input is converted to a meaningful representation.

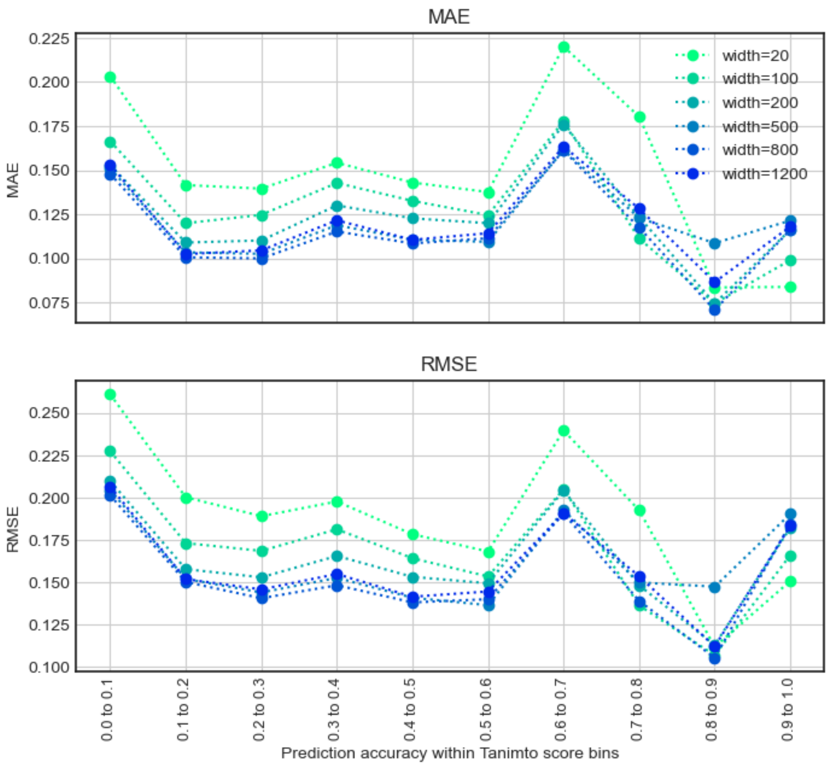


**Figure S7.** Different MS2DeepScore models were trained for different main dense layer widths ranging from 20 to 1200 nodes. The MAEs (upper plot) and RMSEs (lower plot) are here calculated for all spectrum pairs within the validation set (3597 spectra) which fall into one of the 10 possible Tanimoto score bins (x-axis labels).

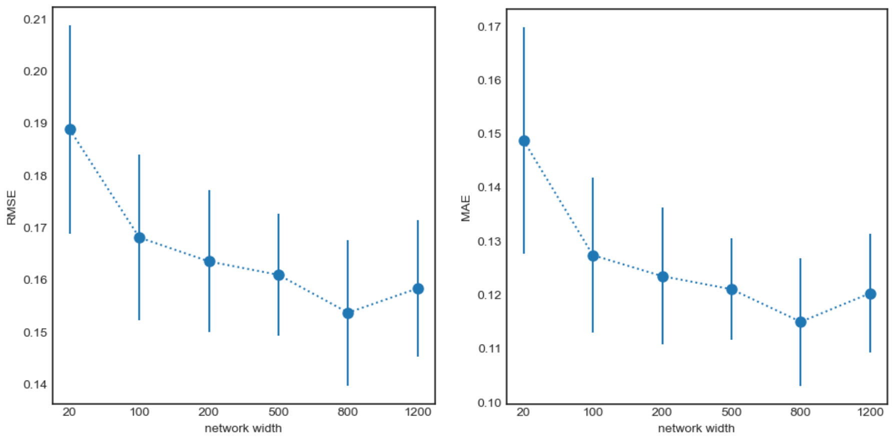


**Figure S8.** Plot showing the mean values for the RMSEs (left) and MAEs (right) for MS2DeepScore models of central network widths between 20 and 1200 nodes. Means are simply the means across all 10 Tanimoto score bins shown in figure (figure S7). Error bars range from -0.5STD to +0.5STD based on the RMSE for all 10 Tanimoto bins.

### Early stopping

We applied early stopping during model training to avoid overfitting. Model training was stopped once the loss on the validation set did not improve for 5 epochs in a row.

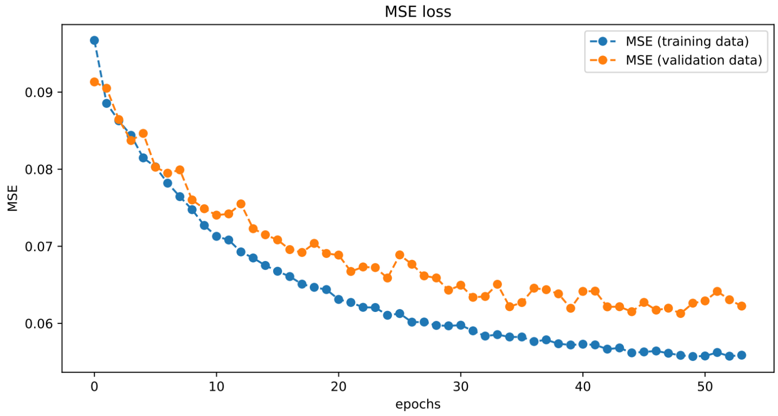


**Figure S9.** Training and validation loss during training of the model.
